# Improved 7 Tesla Transmit Field Homogeneity with Reduced Electromagnetic Power Deposition Using Coupled Tic Tac Toe Antennas

**DOI:** 10.1101/2020.11.06.371328

**Authors:** Tales Santini, Sossena Wood, Narayanan Krishnamurthy, Tiago Martins, Howard J. Aizenstein, Tamer S. Ibrahim

**Affiliations:** Department of Bioengineering, University of Pittsburgh, Pittsburgh, PA, United States of America; Department of Psychiatry, University of Pittsburgh, Pittsburgh, PA, United States of America; Department of Radiology, University of Pittsburgh, Pittsburgh, PA, United States of America

## Abstract

Recently cleared by the FDA, 7 Tesla (7T) MRI is a rapidly growing technology that can provide higher resolution and enhanced contrast in human MRI images. However, the increased operational frequency (~297 MHz) hinders its full potential since it causes inhomogeneities in the images and increases the power deposition in the tissues. This work describes the optimization of an innovative radiofrequency (RF) head coil coupled design, named Tic Tac Toe, currently used in large scale human MRI scanning at 7T; to date, this device was used in more than 1,300 patient/volunteer neuro 7T MRI scans. Electromagnetic simulations were performed for each of the coil’s antennas using the finite-difference time-domain method. Numerical optimizations were used to combine the calculated electromagnetic fields produced by these antennas, based on the superposition principle, and successfully produced homogeneous magnetic field distributions at low levels of power deposition in the tissues. The simulations were then successfully validated in-vivo using the Tic Tac Toe RF head coil system on a 7T MRI scanner.

## Introduction

**<u>Magnetic Resonance Imaging (MRI)</u>** is excellent for soft tissue imaging and determination of its metabolites. This technology provides high-resolution images with several different contrasts and it is widely utilized in clinical settings. The MRI signal increases with higher **<u>static magnetic field strength (B_0_)</u>**. Therefore, advancing from standard clinical scanners – with B_0_ of 1.5 **<u>Tesla (T)</u>** or 3T – to the recent FDA cleared 7T provides a major advantage of increased **<u>signal-to-noise ratio (SNR)</u>**^1^. The enhanced SNR can be used either to increase the resolution of the images or to decrease the scanning time (with the use of higher acceleration factors)^1^. Other advantages of 7T field strength are the higher sensitivity to **<u>blood-oxygen-level-dependent (BOLD)</u>** signal, better venous vasculature conspicuity, enhanced angiography, and improved spectroscopy acquisitions^1^.

The operational frequency for proton imaging at 7T is ~297.2 MHz. When compared to lower field strength, the shorter wavelengths associated with higher operational frequencies can cause spatial inhomogeneities in the **<u>radiofrequency (RF)</u>** fields and reduced skin depths; both which can cause voids or regions of low contrast in the images. The higher operational frequencies and RF inhomogeneities can also lead to a higher average and local **s<u>pecific absorption rate (SAR)</u>**, which can cause temperature rise and potential tissue damage^2^.

The methods developed for improving RF excitation homogeneity can be divided into two main categories. First, multiple pulse sequences and acquisition strategies have been developed to increase the insensitivity of the images to the **<u>circularly polarized component of the RF magnetic field responsible for excitation (B_1_^+^)</u>** field inhomogeneities. These strategies include adiabatic pulses^3,4^, transmit SENSE^5,6^, spoke pulses^7^, and the acquisition of two interleaved modes with TIAMO^8,9^. The second category – and the focus of this work – is related to RF coil design and methodology of operation. This often includes the development of multichannel **<u>transmit RF coil (Tx)</u>** systems, in which the resultant electromagnetic fields can be manipulated with the superposition of the fields generated by each coil element^10^. This technique, known as RF shimming, is accomplished by modifying the phases and amplitudes of the RF field produced by each transmit channel towards specific objectives, usually aiming at increasing the global and/or local B_1_^+^ field homogeneity/intensity and reducing SAR.

Previous works have evaluated the **<u>Tic Tac Toe (TTT)</u>** RF head coil design for 7T MRI. In reference^11^, it was demonstrated that a 16-channel TTT multilevel coil can simultaneously drive up to 4 different eigenmodes; one eigenmode from each of the 4 different physical levels of the coil. Reference^12^ provided a theoretical comparison (based on electromagnetic simulations) of the TTT design with the **<u>transverse electromagnetic (TEM)</u>** resonator, demonstrating an improved transmit field homogeneity and load insensitiveness of the TTT design.

In this work, we describe a methodology for optimizing and operating the TTT transmit coil design for human imaging studies. Constrained numerical optimizations of the 16-channel TTT transmit coil were performed based on **<u>finite-difference time-domain (FDTD)</u>** electromagnetic field simulations while considering the RF power losses in the hardware. Two homogeneous (in terms of B_1_^+^ field) RF shim cases for two different input power and SAR efficiency levels were experimentally implemented on the **<u>single channel (sTx)</u>** mode of a MAGNETOM 7T MRI system using commercially available RF power splitters and phase shifters (coaxial cables). In-vivo B_1_^+^ maps, as well as T2 **<u>SPACE (Sampling Perfection with Application optimized Contrasts using different flip angle Evolution)</u>** and T2 **<u>FLAIR (Fluid Attenuated Inversion Recovery)</u>** sequences were acquired for demonstration purposes. The results demonstrate homogeneous 7T neuro imaging. This RF coil system is currently being used in more than 20 patient/disease-based studies funded by **<u>NIH (National Institutes of Health)</u>**. To date, the current fully implemented version of the RF coil system has helped acquire more than 1,300 neuro in-vivo scans in ongoing human MRI studies^13,14^.

## Methods

### 16-channel Tic Tac Toe RF coil design

The Tx coil is based on the TTT antennas, previously described for foot/ankle^15^, breast^16,17^, and head imaging at 7T. Briefly, one TTT panel is composed of eight square-shape transmission lines elements connected to each other in a Tic Tac Toe fashion. Four of these elements are connected to excitation ports, and the other four elements are used for frequency tuning. The antennas are matched and tuned by varying the length of the copper rods inside the outer struts in these elements (Figure 1d). The Tx coil is composed of four TTT panels positioned around the head, resulting in a 16-channel transmit coil (Figures 1a and 1b). The RF shield is composed of double-sided 4μm-thick copper sheets (Polyflon, Germany). Cuts were added on each side of the copper sheet to reduce eddy currents, as described by Zhao et al^18^. For optimal imaging purposes, the RF coil system incorporates an in-house developed 32-channel **<u>receive (Rx)</u>**-only insert^19^.

**Figure 1:**
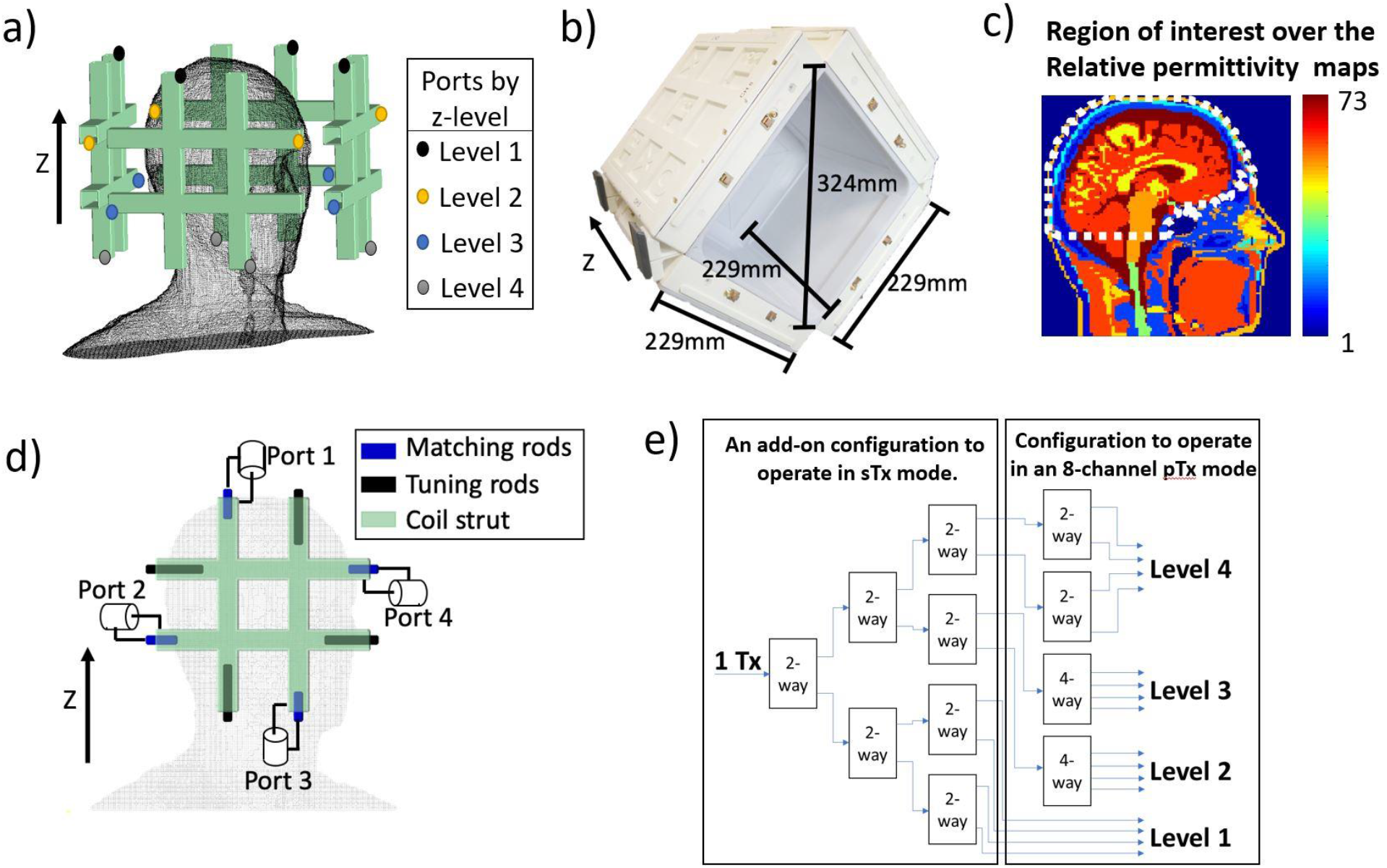
The 16-channel Tic Tac Toe (TTT) Tx head coil FDTD model and experimental implementation. In a), the coil geometry with the location of the 16 channels of the transmit head coil, which are divided into 4 levels in Z-direction; In b), the assembled 16-channel TTT Tx coil and its dimensions; In c), the region of interest (white dashed line) plotted over the relative permittivity map of the Duke model at ~297.2 MHz (7T proton frequency). In d), the ports, matching rods, and tuning rods of a representative panel of the coil. In e), the RF power splitting configuration - exclusively using 2-way and 4-way Wilkinson power dividers - to drive the 16- channel transmit coil using the system’s sTx and pTx modes. The Tx channels associated with the coil’s levels 1, 2, 3, and 4 (shown in b)) experience normalized voltage amplitudes equal to 1, 0.5, 0.5, and 1/√2, respectively.

### FDTD simulations of the Tic Tac Toe coil

The B_1_^+^ and electrical fields were simulated using an in-house developed full-wave FDTD software with an transmission line algorithm for modeling the RF excitation and the coupling^15,17,20–22^. The spatial resolution utilized for the combined RF coil and load was 1.59 mm isotropic and the temporal resolution was ~3ps. The coil geometry (Figure 1a) was created using MATLAB (MathWorks, USA), totaling 257×257×276 or ~18 million Yee cells. Perfect matching layers were implemented to absorb the irradiating fields, being 8 layers added on the top of the model (towards the Z direction), 12 on the sides, and 32 on the bottom^23^. Each port/channel was excited individually with a differentiated Gaussian pulse, while all the other ports were terminated with a 50Ω load using the transmission line numerical model. The Virtual Family Duke model (version 1.0) was used as the load. The model includes the whole head, neck and the top of the shoulders, totaling 23 different tissues. The **<u>region of interest (ROI)</u>** used includes the head regions from the top of the head through the bottom of the cerebellum and excludes the nasal cavities and ears. The lower ~1 cm of the cerebellum volume is excluded as it contains a minimal number of pixels with brain tissues in the Duke model. The contour of the ROI mask applied over the permittivity map is shown in Figure 1c.

The resolution of the FDTD calculated B_1_^+^ fields was then reduced by a factor of 2 to speed up the RF shimming numerical optimizations (described in the next sections), while the resolution of the electric fields was not changed.

### Strategy for RF shimming

Numerical optimizations designed to minimize specific cost functions were used to manipulate the phases of the RF fields generated by the Tx coil channels. When the optimization goal is to increase B_1_^+^ homogeneity, the **<u>coefficient of variation of the B_1_^+^ field</u>** 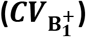 is commonly used as the cost function; however, it may produce local regions of high or low flip angles^24^. Moreover, even if the 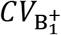 is in its global minimum, SAR levels could be elevated. To overcome these challenges, the cost functions utilized in this work were combinations of 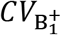, **<u>maximum B_1_^+^</u>** 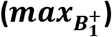, **<u>minimum B_1_^+^</u>** 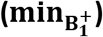, and average SAR. The 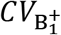 function is continuous and inherently smooth over the multi-dimensional space^25^. As a result, there is a reduced number of local minima produced by this function, which improves the convergence rate for gradient descending algorithms. For this reason, we firstly conducted optimizations using the 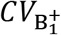 as the cost function and using random phases as the initial conditions (the amplitude values were fixed, following the strategy defined in Figure 1e). The first optimization results were used as the initial conditions for the subsequent optimization, utilizing other cost functions.

The cost functions were minimized using the MATLAB function *fmincom*, including GPU acceleration for the objective and cost functions calculation. The algorithm used for the 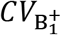 optimizations was the active-set, due to its high efficiency and high convergence rate from random initial conditions^26^. For the optimizations using other cost functions, the interior-point algorithm was used in combination with the initial phase conditions attained from the 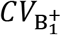 optimizations, since the interior-point algorithm usually performs small steps around the initial conditions without the use of large steps which would dramatically affect the results.

There is an inverse relationship between 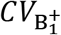 and B_1_^+^ efficiency in most multichannel Tx coil designs, since a highly homogeneous B_1_^+^ can be achieved at the expense of very low efficiency^27^. In this scenario, a high input voltage is required to achieve the desired flip angle, potentially reaching or exceeding the SAR limits and hardware capabilities. To ensure an adequate efficiency for the Tx coil, the optimizations in this work were performed constraining the mean 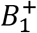 field to 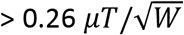 in the sTx mode. This guarantees that there is enough B_1_^+^ intensity to produce 180 degrees flip angle on average with 1ms square pulse and using 8kW RF power amplifier. This constraint also considers the losses between the RF power amplifier and the RF coil plug as well as the losses associated with the coil structure, coil ports, coil plugs, cables, connectors, and power splitters.

### RF shimming for the Tic Tac Toe coil

#### Step 1: RF power splitting

The TTT RF coil system was designed to work in either the sTx or **<u>parallel-transmit (pTx)</u>** modes with the use of Wilkinson power splitters (Figure 1e). Based on the eigenmodes of the Tx coil design^11^, the 16-channel Tx coil is composed of 4 excitation levels spatially positioned in the Z direction (Figure 1a); each level is composed of 4 Tx coil channels. In this work, Level 1 was chosen to be excited with half of the total supplied RF power because of the higher power efficiency associated with this level and its capacity to produce center bright, as previously demonstrated in reference^11^. Levels 2 and 3 play an essential role in exciting the lower brain regions, including the hippocampus and the temporal lobes^11^. One-eighth of the total supplied RF power was used to excite each of these two levels. Level 4, which produces a similar B_1_^+^ pattern of Level 1^11^, received one-quarter of the total supplied RF power. In order to compare this arrangement with other possible configurations, RF shimming optimizations were performed by randomly permutating the Tx coil channels amplitudes with the four possible values of amplitudes, as described in Figure 1e, and then performing phase-only optimizations as shown in Figure 3c.

#### Step 2: Optimization of the B_1_^+^ field and SAR for a specific RF power splitting configuration

SAR calculation was incorporated into the optimization software by sampling the electric fields (sampled random points every 4×4×4 voxels) and calculating the average SAR in these points. This method represents a good estimation of the global average SAR and was used to speed up the optimizations. It is worth noting that this method was only applied inside the iterations of the optimization software, the final SAR was calculated according to the 10g averaging guidelines^28^, using the whole electric fields. The objective function in Equation 1 was minimized in multiple optimization runs, aiming at achieving a compromise between B1^+^ homogeneity and SAR reduction.

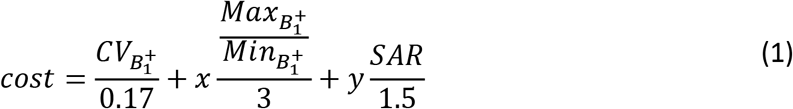

where 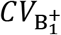, 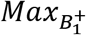, 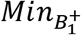 are, respectively, the coefficient of variation, the maximum, and the minimum of the 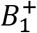 fields inside the region of interest; the constants 0.17, 3, and 1.5 were used to normalize the data and are roughly the expected values of the 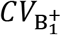,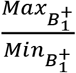 and SAR, respectively; the unit of *SAR* is W/Kg for 2*μ*T; x and y are the constants randomly varying in every optimization in order to change the weights of 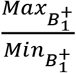 and SAR, respectively, in the cost function.

### Experimental implementation

Two RF shim cases were experimentally implemented using coaxial cables as fixed phase shifters and Wilkinson power splitters (Figure 1e). The imaging experiments were conducted in a whole body 7T scanner (MAGNETOM, SIEMENS, Germany) with 8kW power amplifier capabilities. In-vivo images were acquired in healthy volunteers with informed consent as part of an approved study by the University of Pittsburgh’s Institutional Review Board (identification number PRO17030036). All procedures complied with relevant guidelines and regulations for investigational use of the device in humans.

B_1_^+^ maps were acquired using the Turbo-FLASH sequence^29^ with the following parameters: TR/TE = 2000/1.16 ms; flip angle from 0° to 90° in 18 degrees increments; acquisition time = 12 min, resolution 3.2 mm isotropic. The output was fitted to a cosine function to produce the flip angle maps. For demonstration purposes, 2D FLAIR and 3D T2-SPACE sequences were used to acquire whole-brain images. The respective sequences parameters were: 1) 2D FLAIR, TE/TI/TR = 103/2,900/13,500 ms, BW = 230 Hz/pixel, resolution 0.7×0.7×2 mm^3^, acceleration factor 2, 4 interleaved acquisitions (64 transversal slices), total acquisition time = 7:36 min; 2) 3D T2-SPACE sequence (WIP692), TE/TR = 369/3,400 ms, BW 488 Hz/pixel, T1/T2 = 1,500/250 ms, acceleration factor 3, 224 slices in transversal acquisition, acquisition time = 8:11min, and the receive profile removed using the SPM software.

## Results

Figures 2a and 2b compares the measured and simulated scattering parameters of the 16-channel TTT Tx coil. The simulated and measured values have Pearson correlation coefficient of 0.935. The measured coupling between the opposite ports of each TTT panel (highest coupling of the design) was −4.26dB on an average while the simulated value was −2.68dB on average. This difference is attributed to losses in the copper, elements, and ports of the Tx coil, which are not included as part of the FDTD modeling. The measured maximum and average of coupling between any pair of panels were −17.0dB and −24.4dB, respectively.

**Figure 2:**
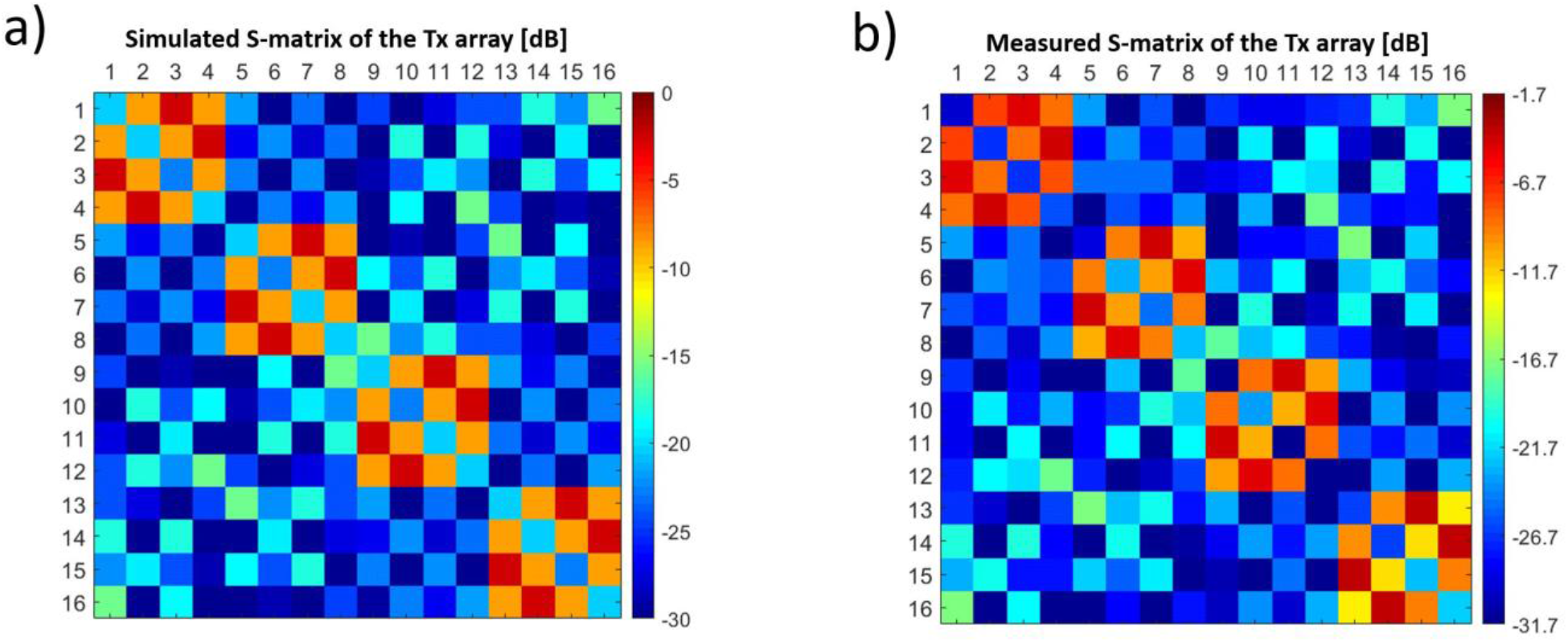
S-parameter comparison between simulations and experiments of the Tic Tac Toe 16-channel Tx head coil. In a), the FDTD simulated s-matrix using a transmission line model mechanism; in b), the experimentally measured s-matrix of the constructed head coil. The color scale limits were modified to compensate for the losses in the constructed coil (not included in the simulations).

Figures 3a and 3b show the effect of permutating the values of the amplitudes associated with each individual Tx channel on the overall 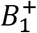 performance. The RF power splitting scheme shown in Figure 1e is compared with approximately 300,000 permutations of the amplitude values followed by phase-only RF shimming. The results – utilizing the RF power splitting scheme from the eigenmodes of the RF coil (figure 1e)– demonstrate superior performance in terms of 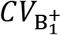, 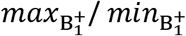, and 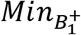. The RF shim case indicated by the black arrow in Figures 3a and 3b was chosen as the starting point for the next step in the optimizations, utilizing the cost function shown in Equation 1.

**Figure 3:**
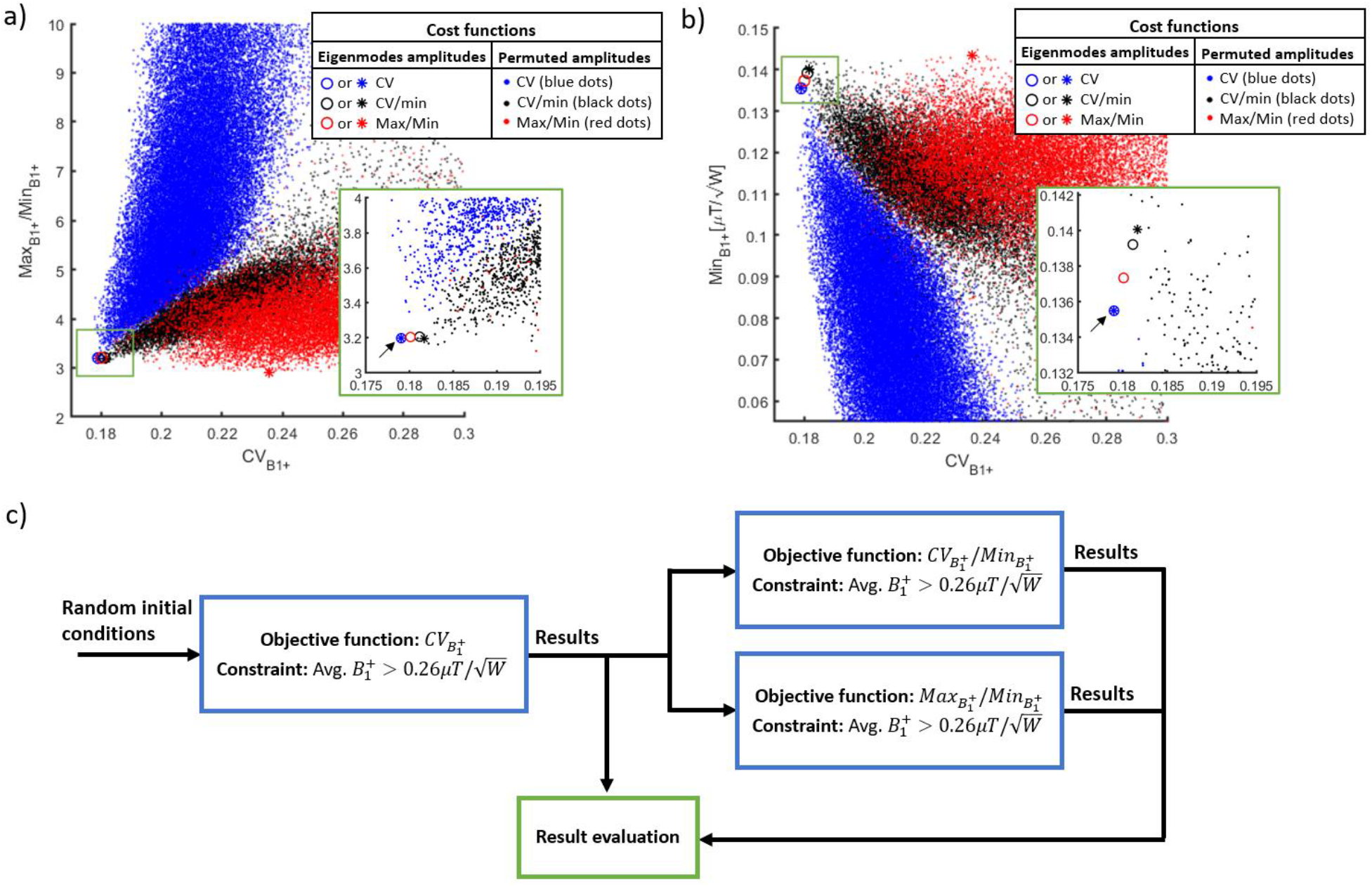
Phase-only B_1_^+^ RF shim cases for the 16-channel Tic Tac Toe RF coil. This analysis investigates the performance of several configurations using 2-way and 4-way splitters for implementation on the sTx mode. In a) and b), the phase-only RF shim cases, with the amplitude scheme derived from the eigenmodes of the RF coil (described in Figure 1b) and in reference^11^, were compared with RF shimming optimizations where the amplitudes of the Tx channels were randomly permutated but can only take on normalized values = 1, 1/√2, or 0.5. Approximately 300,000 optimizations were performed, presented as the colored dots. The cost functions for the RF shimming optimizations were the 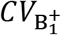, 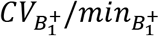, and 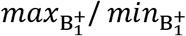 following the flowchart in c). The region of interest for the B_1_^+^ field stats is the entire head from cerebellum excluding the nasal cavities and the ears (Figure 1c). The black arrows point to the case selected as initial condition for the next optimizations, which was chosen due to having a combination of low 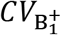, low 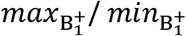, and high 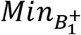. The circles represent the RF shim cases with the best 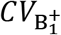 and the asterisks are the cases with the best 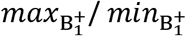.

Figure 4a and 4b show the characteristics of the B_1_^+^ field and SAR variating the weights of the cost function (Equation 1) and variating constraints on the average B_1_^+^ intensity. While higher levels of SAR efficiency (i.e., lower average SAR) and higher power efficiency can be achieved, it usually comes at the cost of lower levels of B_1_^+^ field homogeneity (higher 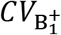 and/or 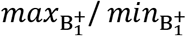). From the scattering plots, we chose two cases for experimental implementation (arrows in Figures 4a and 4b).

**Figure 4:**
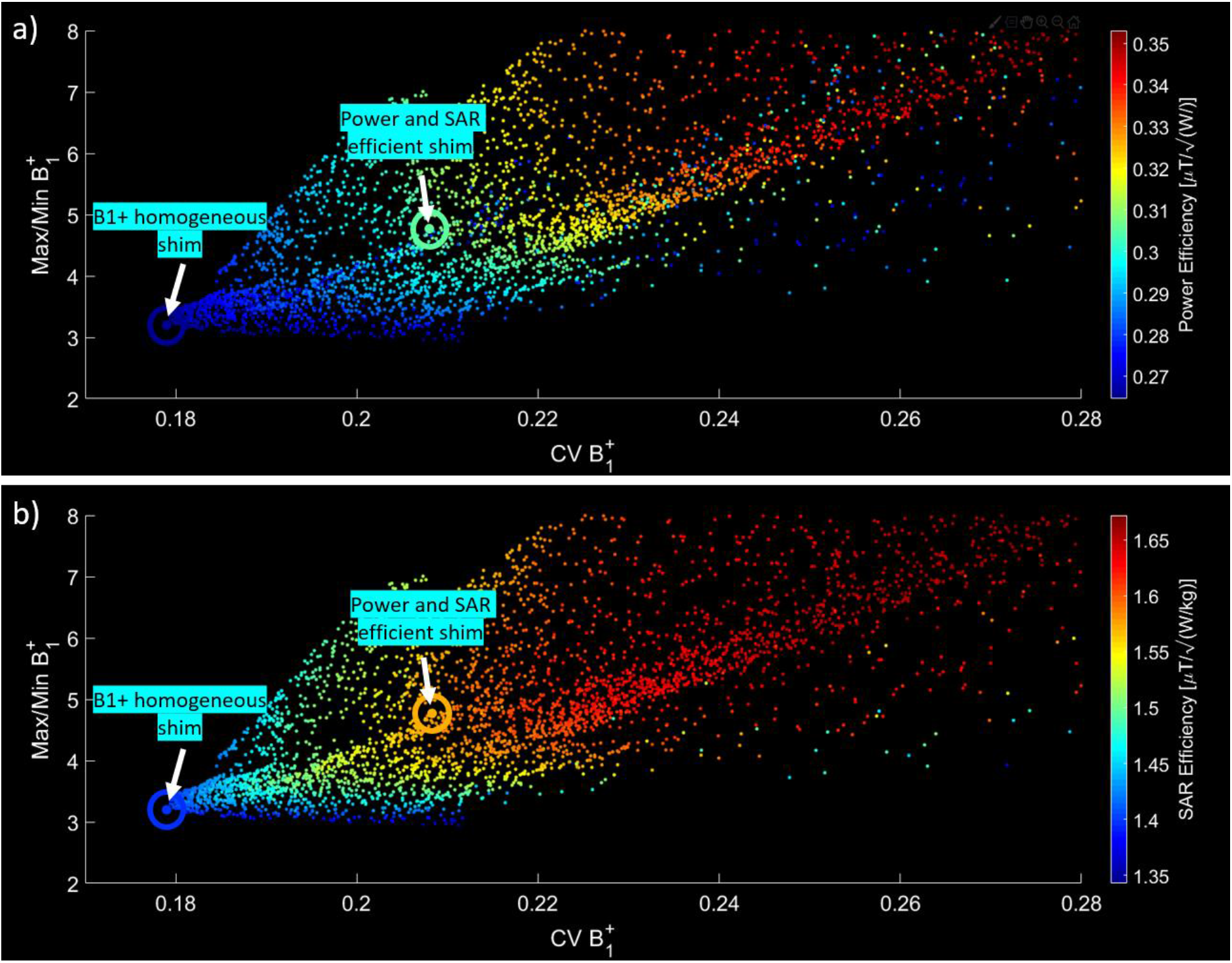
SAR and B_1_^+^ phase-only RF shimming of the 16-channel Tic Tac Toe RF coil. The B_1_^+^ homogeneity parameters (B_1_^+^ CV and B_1_^+^ max/min in x and y axes of the plot, respectively) are compared with the power efficiency (a) and SAR efficiency (b). Each point corresponds to an RF shim case using a cost function (Equation 1) that includes B_1_^+^ CV, B_1_^+^ max/min, average B_1_^+^, and average SAR. Two RF shim cases were chosen for implementation: 1) power and SAR efficient and 2) homogenous B_1_^+^.

Figures 5a-5c show a comparison of the two experimentally implemented, non-subject-specific RF shim cases. Specifically, the comparison shows the B_1_^+^ field maps and FLAIR images acquired on the same volunteer, and the corresponding simulated results on the Duke model. In Figure 5a, a power and SAR efficient RF shim case is shown. It achieves a SAR efficiency of 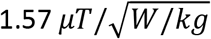 (simulated), average B_1_^+^ of 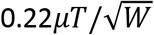 (experimental), 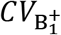 of 20% (experimental), and 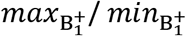 of 4.77 (simulated). The B_1_^+^ homogeneous RF shim case (Figure 5b) has a SAR efficiency of 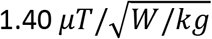 (simulated), average B_1_^+^ of 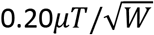 (experimental), 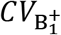 of 17% (experimental), and 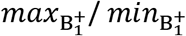 of 3.20 (simulated). Figure 5c shows the profiles in the central slices of the experimental and simulated B_1_^+^ field for the two RF shim cases. The impact of having lower B_1_^+^ field intensities can be seen near the yellow arrows displayed over the B_1_^+^ field distributions and the FLAIR images in Figures 5a and 5b.

**Figure 5:**
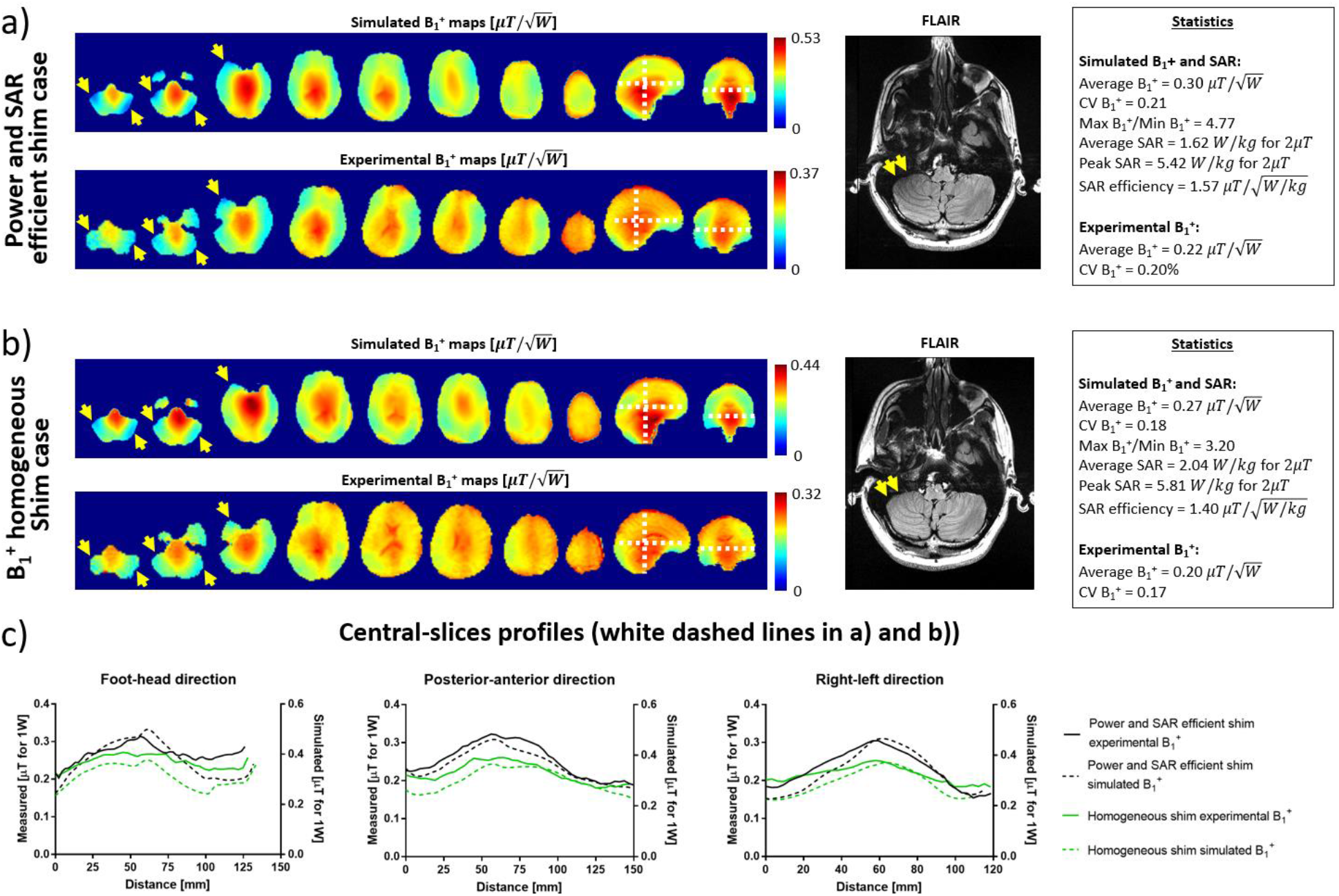
Simulated and experimental B_1_^+^ maps comparisons and in-vivo FLAIR images for 2 RF shim cases. In a), simulated and in-vivo B_1_^+^ maps, FLAIR acquisition, and statistics of the power and SAR efficient RF shim case. The statistics from the simulations were calculated over the region of interest shown in Figure 1c. In b), simulated and in-vivo B_1_^+^ maps and FLAIR acquisition (with same subject as a)) for the B_1_^+^ homogenous RF shim case. The yellow arrows point to regions of low B_1_^+^ that were mitigated with the B_1_ ^+^ homogeneous RF shim case. In c), the central slice profiles from the simulated and experimental B_1_^+^ maps (white dashed lines in a) and b)).

Figure 6 shows representative slices of the T2-weighted 3D SPACE images acquired in a volunteer with large head size (approximately 205 mm in anterior-posterior direction from the forehead) using the B_1_^+^ homogeneous RF shim case shown in Figure 5b. Using an MR sequence that requires high levels of B_1_^+^ homogeneity, the images show that the implemented RF shim case provides non-subject specific field distribution and it achieves full brain coverage – including the cerebellum and temporal lobes – in a relatively larger head.

**Figure 6:**
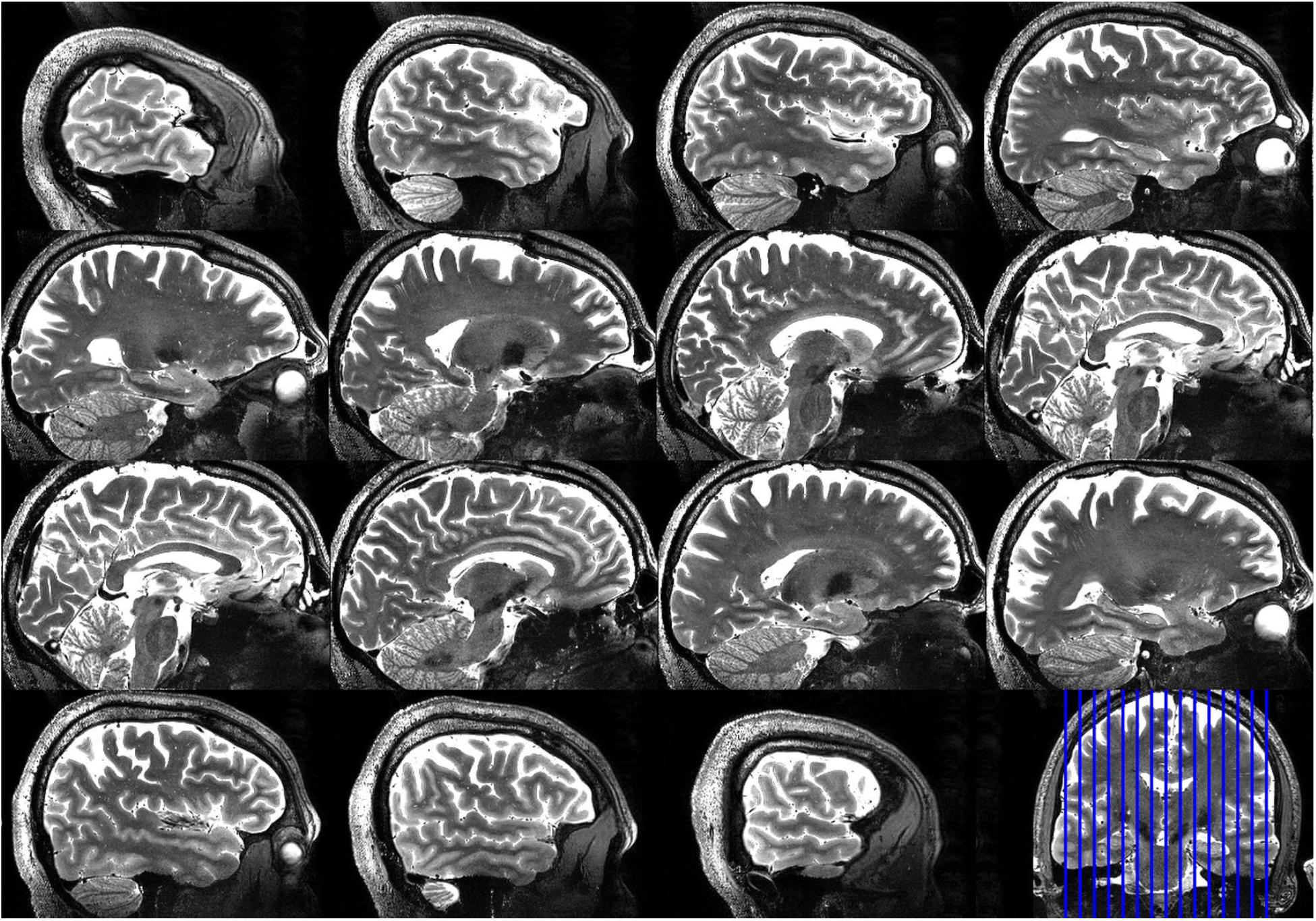
Sagittal slices of the 3D-SPACE acquired at 0.6mm isotropic resolution, showing full brain and cerebellum coverage in a volunteer with large head size (~205mm in anterior-posterior direction measured from the forehead). The images were obtained using the B_1_^+^ homogenous RF shim case on the sTx mode. The parameters of the acquisition were: TE/TR = 367/3400 ms, acceleration factor 3, BW = 434Hz/Px, transversal acquisition of 224 slices, field of view 192×165.6 mm in axial plane, and acquisition time = 8:11 min.

## Discussions

In this work, a 16-channel transmit coil based on the Tic Tac Toe design was optimized using non-subject specific phase-only RF shimming. The RF shimming approach combined with the coil’s 4 excitation levels in the Z direction can significantly impact the Tx coil performance by introducing tradeoffs among RF power efficiency, B_1_^+^ field homogeneity, and SAR. These parameters can be controlled, depending on the imaging application and the RF amplifier capacity, by changing the lengths of the coaxial cables feeding the coil or by using pTx systems.

### S-matrix comparison

The RF TTT transmit coil was simulated with a transmission line algorithm for modeling the excitation and coupling as part of the in-house developed FDTD package^30^. Accurate electromagnetic modeling of the coil’s coupling is critical for implementing RF shimming fully based on B_1_^+^ and electric fields calculated from electromagnetic simulations. The simulated and measured s-matrices are highly comparable (Figures 2a and 2b, respectively), with a Pearson correlation coefficient of 0.935. There is an offset in the values (−1.7dB on average) that is related to the losses in the coil and connectors.

### Strategy for power splitting among the channels

The amplitudes for the Tx channels were chosen based on the eigenmodes of the RF coil so that the most efficient coil levels (demonstrated in reference^11^) are excited with a larger fraction of the supplied RF power. This approach produces improved B_1_^+^ homogeneity (Figure 3a) and high minimum intensity (Figure 3b) when compared with other possible combinations using the same RF power splitter configuration. All cases presented can be easily implemented using 2-way and 4-way RF power splitters and coaxial cables as phase shifters on the system’s sTx mode.

### B_1_^+^ and SAR RF shimming

Figures 4a and 4b show the flexibility of the phase-only RF shimming scheme in combination with the 16-channel TTT coil design, demonstrating the tradeoff among SAR efficiency, B_1_^+^ efficiency, and B_1_^+^ homogeneity. Based on the electromagnetic simulations, a homogeneity of 18% 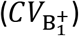 can be achieved within the ROI, resulting in a SAR efficiency of 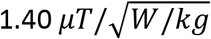 and an average B_1_^+^ of 0.27 *μT*. Increasing the weight of the SAR in the cost function (Equation 1) and increasing the mean B_1_^+^ constraint improved the SAR efficiency to 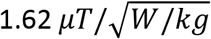 while maintaining a 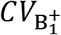 of 21% and an average B_1_^+^ field of 0.30 *μT*. These values of SAR efficiency and B_1_^+^ field homogeneity represent an improved performance for 7T RF head coils when compared with traditional RF coils. As a comparison, Krishnamurthy et al.^12^ reported the values of SAR efficiency of 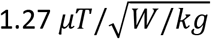 and homogeneity of 27% for the TEM resonator, using the same simulation environment utilized in this work. Other studies investigated the SAR performance for several other coil designs^31–34^, however the simulations were performed in different software environments and often using different head models, which makes the comparison with this work challenging. The ROI (Figure 1c) was chosen so that the RF fields become more consistent with different brain sizes, when compared with a specific brain size (in the case of a brain-only ROI).

In general, the cost function combining 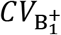, 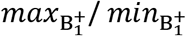, and *SAR* with the average 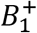 constraint presented a good performance for this particular RF coil design, as it allowed different operational points where it is possible to minimize 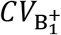, 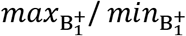, or maximize SAR efficiency (Figures 4a-b), depending on the application. It is important to note that the optimal cost function may vary depending on the RF coil design, the constraints included in the simulation, and the ROI used.

### Experimental verification

Figures 5a and 5b show the experimental implementation of the two RF shim cases and the corresponding simulated B_1_^+^ maps. The first case is intended for higher SAR and higher RF power efficiency; the second case is optimized for higher B_1_^+^ homogeneity. The images - using a high flip angle sequence (FLAIR) - demonstrate the differences between the two RF shim cases in the cerebellum, which is a very challenging region at 7T MRI^35,36^. Although the B_1_^+^ homogeneous RF shim case presents lower RF power efficiency, the improved B_1_^+^ homogeneity justify its use, since the lower RF power efficiency can be compensated by adjusting the input voltage in the coil or by increasing the pulse width of the sequences.

Figure 5c shows a comparison between the simulated and experimental B_1_^+^ profiles. The differences in the intensity can be explained by the losses in the splitters/cables/connectors (approximately ~17% measured loss in voltage) and losses in the coil. The differences in the field distribution can be attributed to the model approximations and differences between the head model size/position and the in-vivo. For instance, the brain in the model used is about 14cm long in foot-head direction, while the volunteer is about 12.5cm.

The 3D T2-SPACE whole-brain and cerebellum acquisition, shown in Figure 6, was acquired at 7T using the B_1_^+^ homogeneous RF shim case implemented on the sTx mode (Figure 5b). The results demonstrate that homogeneous whole-brain imaging in a large head size volunteer is achievable with a challenging T2-weighted sequence at 7T MRI^37–39^ – often associated with significantly lower signal in the temporal lobe and cerebellum.

The RF coil design presented in this work has considerably lower power efficiency when compared to traditional 7T RF head coil designs. This is attributed to the strong coupling between opposite elements of the TTT coil, which causes a significant portion of the transmitted power to be dissipated in the RF power splitters and the scanner system’s circulator. For instance, the two RF shim cases presented in this work have a mean B_1_^+^ of 0.27 and 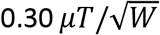 in the simulations (not considering losses in the hardware). Using the same software environment, Narayanan et al.^12^ demonstrated that the 4-channel, 16-element TEM resonator (with the same length as the 16-channel TTT coil) presents a power efficiency of 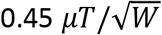. However, considering all the power losses in the system and in the coil, the 16-channel TTT coil still provides enough mean B_1_^+^ intensity to have inversion using 1ms square pulse with 8kW power amplifier (standard in older 7T scanners), which is sufficient for most imaging applications. The B_1_^+^ homogeneous RF shim case (Figure 5b) is currently being used with the sTx mode on more than 20 NIH patient/disease studies conducted in our facility^13,14^ due to its high B_1_^+^ homogeneity and extended coverage in challenging-to-image regions in the head at 7T.

## Data availability

The data that support the findings of this study are available from the corresponding author upon request.

## Acknowledgments

This work was supported by NIH R01MH111265, R01AG063525, and R01EB009848. The author Tales Santini was partially supported by the CAPES Foundation, Ministry of Education of Brazil, 13385/13-5. This research was also supported in part by the University of Pittsburgh Center for Research Computing (CRC) through the resources provided.

## Author contributions

Conceptualization: TS, HJA, TSI; Simulation of the coil design: NK, TS; Implementation of the optimization methods: TS, SW; MRI image acquisition: TS, TM, TSI; Initial manuscript draft: TS, TSI; Manuscript revision: all authors.

## Additional Information

**Competing financial interest**: the authors declare no competing financial interest.

